# ELF3 polyQ variation in *Arabidopsis thaliana* reveals PIF4-independent role in thermoresponsive flowering

**DOI:** 10.1101/038257

**Authors:** Maximilian O. Press, Amy Lanctot, Christine Queitsch

## Abstract

Plants have evolved elaborate mechanisms controlling developmental responses to environmental stimuli. A particularly important stimulus is temperature. Previous work has identified the interplay of PIF4 and ELF3 as a central circuit underlying thermal responses in *Arabidopsis thaliana*. However, thermal responses vary widely among strains, possibly offering mechanistic insights into the wiring of this circuit. ELF3 contains a polyglutamine (polyQ) tract that is crucial for ELF3 function and varies in length across strains. Here, we use transgenic analysis to test the hypothesis that natural polyQ variation in ELF3 is associated with the observed natural variation in thermomorphogenesis. We found little evidence that the polyQ tract plays a specific role in thermal responses beyond modulating general ELF3 function. Instead, we made the serendipitous discovery that ELF3 plays a crucial, PIF4-independent role in thermoresponsive flowering under conditions more likely to reflect field conditions. We present evidence that ELF3 acts through the photoperiodic pathway, pointing to a previously unknown symmetry between low and high ambient temperature responses. Moreover, in analyzing two strain backgrounds with vastly different thermal responses, we demonstrate that responses may be shifted rather than fundamentally rewired across strains. Our findings tie together disparate observations into a coherent framework in which multiple pathways converge in accelerating flowering in response to temperature, with some such pathways modulated by photoperiod.

## INTRODUCTION

The responses of plants to temperature variation are of central importance to food security in a changing world [1]. Therefore, the elucidation of the genetic pathways underlying these responses has been a key mission of plant science [2]. Many previous studies examined the phenomena of circadian temperature compensation [3–5], thermoresponsive flowering [6–10], and temperature effects on plant morphology [11–16]. Several have converged on PIF4 as a master regulator of temperature responses, and ELF3 as an input to PIF4 integration, among many other genes and pathways [14–16]. Given known regulatory interactions between ELF3 and PIF4 [17–20], it is reasonable to predict that both operate in the same pathway for thermal response phenotypes [21]. Recent reports focusing on one such phenotype, hypocotyl elongation, support this expectation [14–16].

ELF3 serves to repress hypocotyl elongation by reducing PIF4 levels. This repression of PIF4 occurs at both the transcriptional level, through the role of ELF3 in the Evening Complex (EC) [17,19], and at the post-translational level, through PIF4 destabilization by phytochrome phyB [22]. Light sensing enforces circadian oscillations of the EC and other components, leading to calibration of the circadian clock [23,24], resulting in diurnal repression of hypocotyl elongation through repression of PIF4 and PIF5 [17,19]. ELF3 also plays a crucial role as a flowering repressor [25]. Consequently, *elf3* null mutants show elongated hypocotyls even in the light, and flower early.

PIF4 is one of a family of basic helix-loop-helix (bHLH) “phytochrome-interacting factors” (PIFs), transcription factors with overlapping functions promoting skotomorphogenesis. Under dark conditions, the PIFs act to target phyB for ubiquitin-mediated degradation by the E3 ubiquitin ligase COP1, thereby repressing photomorphogenesis [26]. Under light conditions, degradation of PIFs is mediated by direct interactions with photoactivated phyB [22]. PIF4 is distinct from the other PIFs in having specific roles in temperature sensing and flowering [27]. *pif4* null mutants show short hypocotyls with photomorphogenic attributes even in the dark [28].

At elevated ambient temperatures (27°-29°) the wiring of these signaling pathways changes. Several independent studies have recently found that elevated temperatures, specifically during dark periods [29], inhibit the activity of the EC by an unknown mechanism [14–16], leading to increased expression of *PIF4* and its targets [11,27]. This increased PIF4 activity leads to several morphological temperature responses through various signaling pathways [13,27]. PIF4 is also required for the acceleration of flowering at 27°C under short photoperiods [9, 29], though these observations have been disputed [30,31]. While PIF4 action alone (among PIFs) is essentially sufficient for most described thermomorphogenic responses [11,27], there is evidence for a limited role of PIF5 (though not other PIFs) in thermoresponsive flowering under short days (SDs) [30,31]. In contrast, under continuous light, *pif4* null mutants have an intact temperature-dependent acceleration of flowering [11]. Lastly, *pif4* null mutants lose the normal elongation of petioles under high temperatures [11]. It is unclear why PIF4 does not affect thermoresponsive flowering under continuous light; yet, this phenomenon may reflect low PIF4 levels under these conditions due to inhibition by phyB. Under longer photoperiods and higher temperature a flowering acceleration still exists [7,11], which suggests a PIF4-independent thermoresponsive flowering pathway. Nonetheless, recent reviews of the literature tend to emphasize the primacy of PIF4 in this response [10,32,33], although the condition of elevated temperature with short photoperiods is probably rare in the field.

Recent studies have identified ELF3 as a plausible upstream regulator of PIF4 in thermal responses [14– 18]. However, others have implicated different candidates, such as FCA [13], and mathematical modeling has suggested that ELF3/EC complex regulation alone is insufficient to explain PIF4 thermal regulation [14,34]. The exact mechanisms of this response have yet to be unraveled.

Specifically, the mechanism by which EC/ELF3 activity is reduced under elevated temperatures (“temperature sensing”) is not known. We recently used transgenic experiments to demonstrate that ELF3 function is dependent on the unit copy number of its C-terminal polyglutamine (polyQ) tract [35]. This domain is likely disordered, and disordered domains evince structural changes in response to physical parameters such as temperature [36]. Thermal remodeling of this polyQ tract is a plausible mechanism by which ELF3 activity could be modulated through temperature. This polyQ tract also shows substantial natural variation [35], potentially serving as a factor underlying natural variation in thermoresponsive phenotypes. For example, in flies, variable repeats are associated with local temperature compensation adaptations [37]. In short, the ELF3-polyQ is an attractive candidate for adaptive variation in the ecologically relevant trait of temperature response [38].

In this study, we used transgenic polyQ variants of ELF3 in two *A. thaliana* genetic backgrounds to dissect the contribution of the polyQ tract to temperature response. We show that polyQ repeat copy number modulates temperature sensing by affecting overall ELF3 function. Surprisingly, we found that ELF3’s role in thermoresponsive flowering appears to be entirely independent of PIF4. We postulate that ELF3’s primary role in thermoresponsive flowering is PIF4-independent and occurs through the photoperiodic pathway, and that this role is in turn dependent on the genetic background.

## RESULTS

### The hypocotyl elongation temperature response is modulated by the ELF3 polyQ tract affecting overall gene function

Many recent studies noted the involvement of ELF3 in temperature-dependent hypocotyl elongation [14–16,39], concluding that ELF3 protein activity is reduced under elevated temperatures, thereby relieving ELF3 repression of *PIF4*. *PIF4* up-regulation then leads to the observed hypocotyl elongation. We examined whether polyQ tract variation in ELF3 in two backgrounds affects hypocotyl elongation at 27°C (Fig. 1). We previously showed that ELF3 polyQ variation has pleiotropic background-dependent effects, with nonlinear associations between polyQ tract length and quantitative phenotypes (including hypocotyl elongation at 22°C; ref. 33). Certain variants (16Q for Ws, >20Q for Col) generally complemented *elf3* null mutant phenotypes in Col and Ws *A. thaliana* strains, whereas other variants complemented only specific phenotypes or behaved as hypomorphs across all tested phenotypes. Here, we observed similar trends for thermoresponsive hypocotyl elongation (Fig. 1). For example, in the Ws background (Fig. 1A), the endogenous ELF3 variant (16Q) partially complements the *elf3* null mutant; another variant (9Q) fully complements the hypocotyl temperature response. Other polyQ variants behaved as hypomorphs in Ws. In the Col background (Fig. 1B), the endogenous 7Q variant, among other variants, failed to rescue the response, agreeing with our previous observation that these transgenic lines are hypomorphic in this background [35]. Deleting the entire polyQ tract eliminated thermoresponsive hypocotyl elongation in both Col and Ws backgrounds. We next addressed whether the observed phenotypic variation among polyQ variants was due to variation in thermosensing or variation in general ELF3 function. We found that robust thermal responses were strongly correlated with the overall functionality of each ELF3 variant in hypocotyl elongation (Fig. 1C), such that variants with intact thermal responses exhibited short hypocotyls at 22°C, whereas ELF3 variants with defective thermal responses exhibited elongated hypocotyls regardless of temperature. Furthermore, this ELF3 functionality effect is dependent on genetic background (comparing for instance the 16Q and 20Q responses). Together, these results suggest that the ELF3 polyQ tract controls repression of hypocotyl elongation regardless of temperature, rather than sensing temperature specifically. Nonetheless, our transgenic ELF3 polyQ lines remain informative as an allelic series of ELF3 function to understand the role of ELF3 in the de-repression of PIF4, which is thought to underlie thermomorphogenesis [14–16,39–41].

**Fig. 1.**
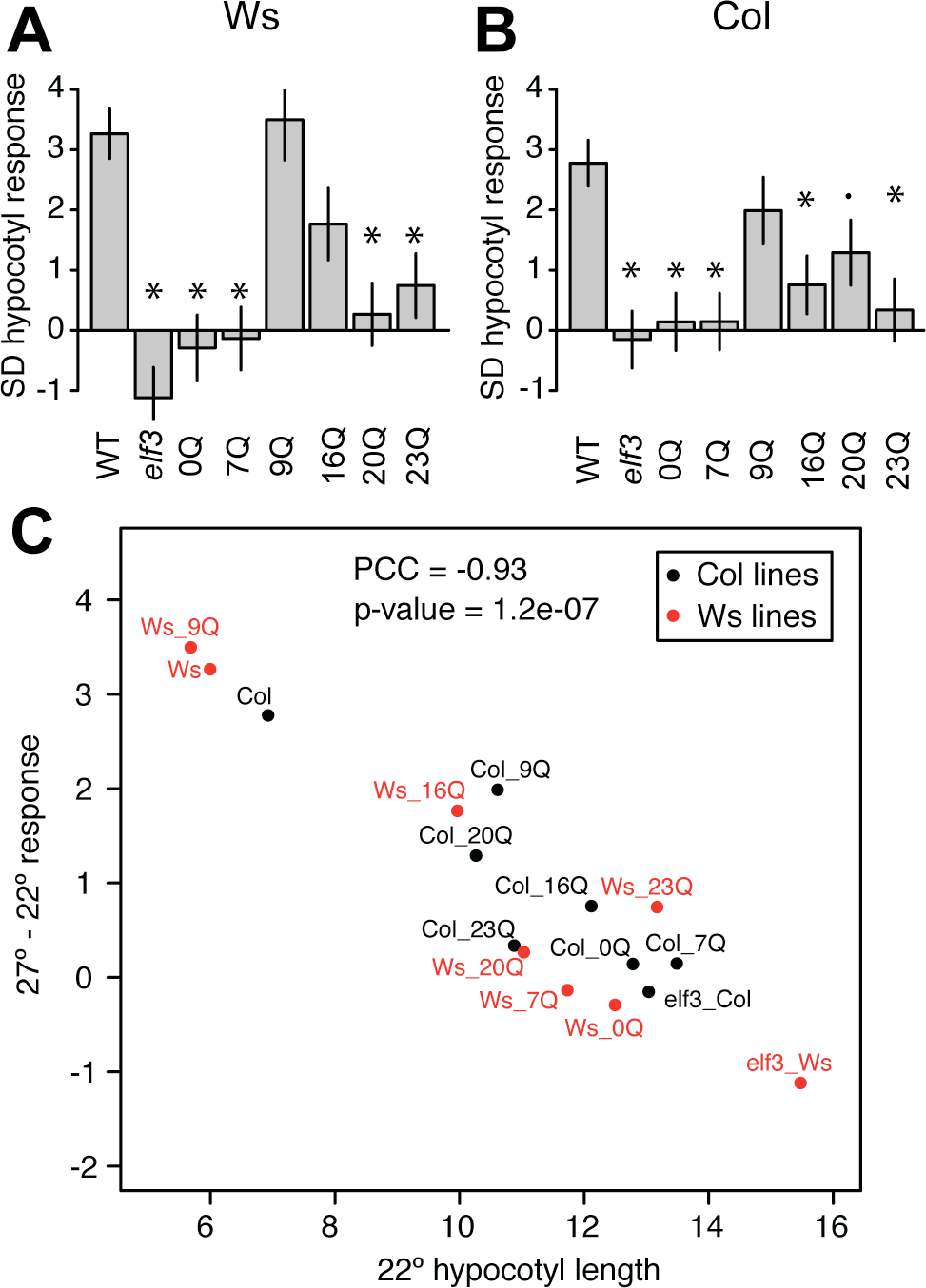
Response to elevated temperature (27°, relative to 22°) among transgenic lines expressing ELF3-polyQ variants. Mean response and error were estimated by regression, based on two independently-generated transgenic lines for each genotype, with n >= 30 seedlings of each genotype in each condition (Table S1). WT = Ws, *elf3*= *elf3* mutant + vector control, 0Q= *elf3* mutant+*ELF3* transgene lacking polyQ, etc. Error bars indicate standard error of the mean. (A): Ws (Wassilewskija) strain background. Lines are generated in an *elf3-4* background. (B): Response in the Col (Columbia) strain background, lines were generated in an *elf3-200* background. In both (A) and (B),**: Bonferroni-corrected p < 0.01, **: Bonferroni-corrected p < 0.05,.: Bonferroni-corrected p < 0.1 in testing the interaction term (different response from WT, Ws or Col). (C): Temperature response is a function of ELF3 functionality (repression of hypocotyl elongation at 22°). Simple means of 22° hypocotyl length, regression estimates of temperature response. PCC = Pearson correlation coefficient; p-value is from a Pearson correlation test.

### Expression of PIF4 and PIF4 targets as a function of temperature and ELF3

To evaluate the hypothesis that the thermal response defects in the transgenic lines was due to up-regulation of PIF4 and PIF4 targets, we measured transcript levels of *PIF4* and its target *AtHB2* in seedlings of selected lines from both backgrounds at 22°C and 27°C (Fig. S1). Like others [15, 16], we observed an inverse relationship between ELF3 functionality and transcript levels of *PIF4* and *AtHB2*, with larger effects on *PIF4* expression. The ELF3 lines with the strongest thermal response (*e.g.* 16Q in the Ws background) showed the most robust de-repression of *PIF4* at elevated temperature. However, *elf3* null mutants retained some *PIF4* up-regulation under these conditions, especially in the Ws background. We conclude that ELF3-mediated de-repression of PIF4 is involved in thermal responses as suggested by prior studies [15, 16]; however, de-repression of PIF4 and its targets may not be sufficient to explain the entirety of thermal response defects in *elf3* null mutants.

### ELF3 polyQ variation affects thermoresponsive adult morphology and flowering time

Following the expectation that ELF3’s thermal response acts through PIF4, we reasoned that ELF3 should also play a role in other PIF4-dependent thermal responses. One well-known response to elevated temperature is adult petiole elongation. *pif4* mutants fail to show this response when grown at elevated temperatures [11]. We measured petiole length in the ELF3 polyQ transgenic lines, expecting that, due to general PIF4 de-repression, poorly-functioning ELF3 polyQ lines would show no response (perhaps due to constitutively elongated petioles, similar to hypocotyls; Fig.2). In stark contrast to this expectation, we found that all lines had a robust petiole response to temperature (Fig. 2A, B). This effect was apparent in both Ws (Fig. 2A) and Col backgrounds (Fig. 2B). Moreover, this response was actually accentuated in *elf3* null mutants and in poorly-functioning ELF3 polyQ variants (Fig. 2A, B).

**Fig. 2.**
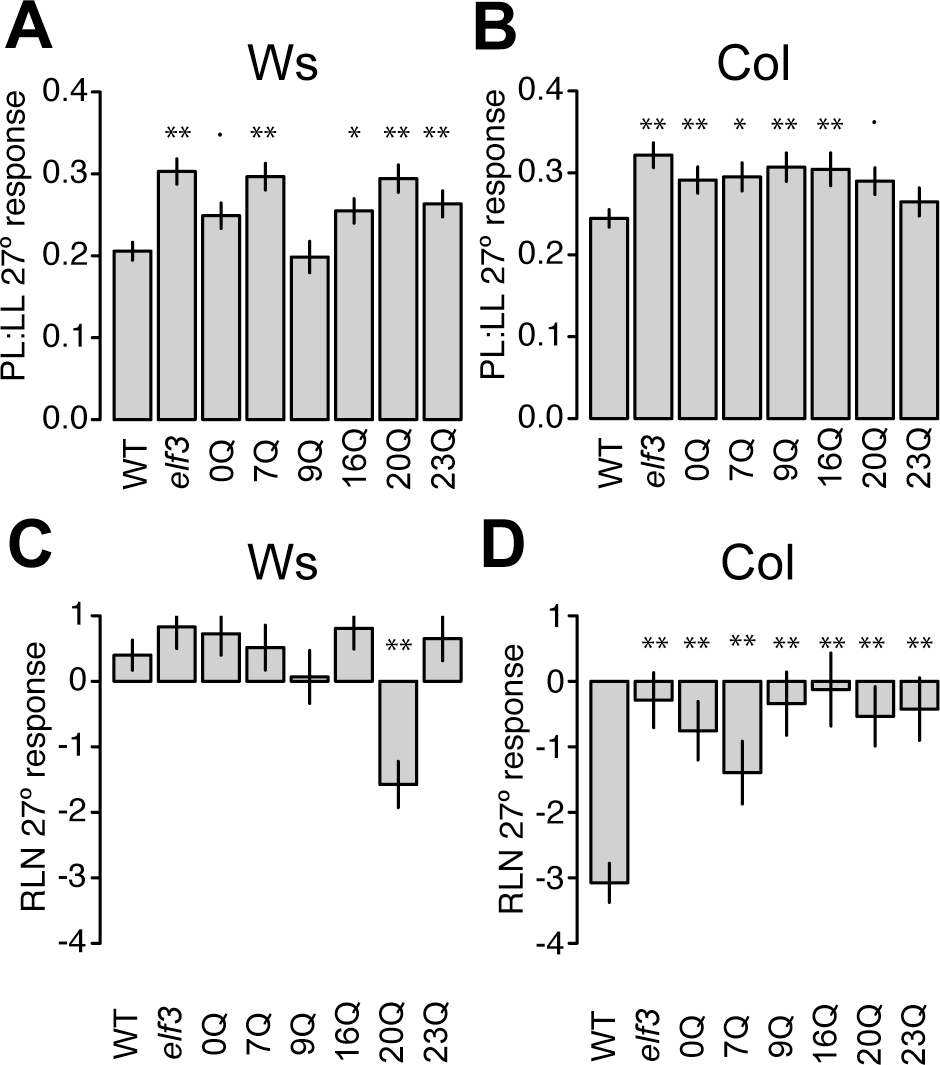
Adult plant responses to elevated temperature (27°, relative to 22°) in long days among transgenic lines expressing different ELF3-polyQ variants. (A) and (C): Response in the Ws (Wassilewskija) strain background. Lines are in an *elf3-4* background. (B) and (D): Response in the Col (Columbia) strain background, lines are in an *elf3-200* background. (A) and (B) display PL:LL temperature response, (C) and (D) display RLN temperature response. Average responses and errors were estimated in a regression model accounting for variation between experiments (Table S2), based on two to three independently-generated transgenic lines for each genotype. n >= 24 plants overall for each genotype in each condition. PL:LL = petiole to leaf length ratio at 25 days post germination, RLN = rosette leaf number at flowering, WT = wild type, *elf3* = *elf3* mutant+vector control, 0Q = *elf3* mutant+*ELF3* transgene with entire polyglutamine removed, etc. Error bars indicate standard error. In each case, **: Bonferroni-corrected p < 0.01, **: Bonferroni-corrected p < 0.05, .: Bonferroni-corrected p < 0.1 in testing the interaction term (different response from WT, Ws or Col).

Further, we measured flowering time in transgenic lines as the number of rosette leaves at flowering (Fig. 2C, D). PIF4 is not required for the accelerated flowering temperature response under longer photoperiods [11]. Hence, we expected that loss of ELF3 function should not affect thermoresponsive flowering if ELF3’s thermal signaling role acts through PIF4. In contrast to this expectation, in the Col background, *elf3* mutants had an abrogated flowering response to elevated temperature (Fig. 2D). Moreover, most variants in the Col background entirely failed to rescue this phenotype, and even the endogenous 7Q shows only a partial rescue. While *elf3* mutants show abrogated flowering response to lower temperatures [8], it was not expected that a similar role would extend to the elevated temperature flowering response, which is usually considered to be dominated by PIF4 [99], gibberellin signaling [30], and other transcriptional regulators of *FT* such as SVP [31].

Unlike Col, Ws lacks a robust flowering response to elevated temperature under these conditions [42], and indeed, variants in the Ws background generally showed no thermoresponsive flowering (Fig. 2C). Thus, ELF3 polyQ variation does not suffice to enhance the negligible thermoresponsive flowering in the Ws background under these conditions. In light of this data, the roles of ELF3 and PIF4 in the elevated temperature response appear to be independent of one another under these experimental conditions and for these traits. These results are intriguing, given that the PIF4 pathway is the best-recognized mechanism for thermoresponsive flowering at high temperatures [9,10,32,33]. Therefore, we suggest that ELF3 acts in a PIF4-independent pathway for thermoresponsive flowering at high temperatures.

### ELF3 regulates thermoresponsive flowering under long days, and is not required for PIF4-dependent adult thermomorphogenesis

Our results with ELF3-polyQ variants suggested that ELF3 dysfunction does not meaningfully affect PIF4-dependent traits, but does affect PIF4-independent traits in adult plants in long days. However, these results may be due to subtle differences in conditions between our approach and those used by previous investigators. We therefore directly addressed the relationship of ELF3 and PIF4 in adult thermoresponsive phenotypes by growing *pif4* and *elf3* mutants with various thermal treatments. Previous experiments with *pif4* mutants used different conditions from ours, specifically a later transfer to elevated temperature [11]. Hence, it was possible that the observed inconsistencies between *elf3* and *pif4* effects on adult thermoresponsive phenotypes were a trivial consequence of experimental conditions. Specifically, the effects of elevated temperature during the early seedling stages (the conditions we use) may induce pathways irrelevant to treatments at later, vegetative stages. Thus, we tested both transfer conditions under long days (Fig. 3). We found that the effect of different experimental conditions is negligible, though the earlier 27°C treatmentshowed a slightly stronger morphological response (Fig. 3A, B). Thus, the timing of the 27°C treatment (early seedling vs. vegetative stage) does not substantially affect adult thermoresponsive traits. Further, our results under long days were similar to previous observations under continuous light [11], showing that PIF4 is essential for petioleelongation (Fig. 3B), but dispensable for thermoresponsive flowering (Fig. 3C). Our PIF4 results were in direct contrast to ELF3, which was dispensable for petiole elongation (Fig. 3B), but essential for thermoresponsive flowering (Fig. 3C). These results indicate the apparent independence of ELF3 and PIF4 in these specific responses, and suggest that seedling thermomorphogenesis, adult thermomorphogenesis, and thermoresponsive flowering constitute three independent developmental responses.

**Fig. 3.**
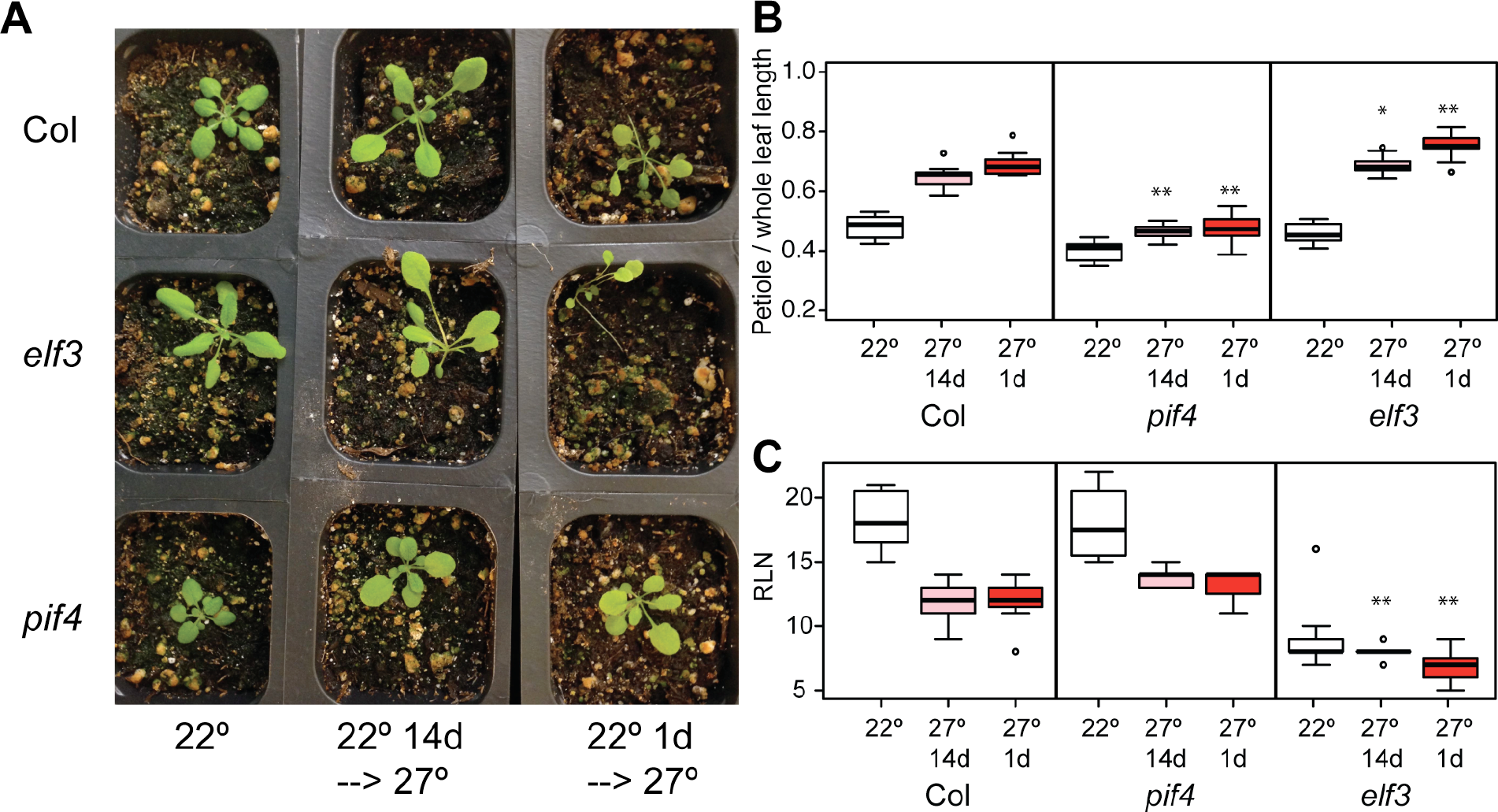
*elf3* and *pif4* null mutant phenotypes are independent under LD treatments and robust to conditions. (A), (B), and (C): 22°:constant 22° LD growth; 27° 14d: transfer from 22° to 27° at 14 days post-germination; 27° 1d: transfer from 22° to 27° at 1 day post-germination. (A): Col (WT), *elf3-200*, and *pif4-2* plants grown under long days with three different temperature regimes were photographed at 20 days post germination. Experiment was repeated with similar results. (B and D): Petiole elongation responses of the indicated genotypes, measured by ratio of petiole to whole leaf length at 25 days post germination. Regression analysis of data in Table S3. In each case, **: Bonferroni-corrected p < 0.01, **: Bonferroni-corrected p < 0.05, .: Bonferroni-corrected p < 0.1 in testing the interaction term (different response from Col).

One open question was whether the dispensability of ELF3 for petiole elongation reflected increased importance of other inputs to PIF4, such as FCA, which is involved in PIF4-dependent thermoresponsive petiole elongation in 7-day-old seedlings [13]. We therefore measured adult thermoresponsive petiole elongation in *fca* mutants (Fig.S2A), and unexpectedly found no substantial difference between *fca* mutants and WT Col. Regulatory rewiring across development may remove FCA and ELF3 as inputs to PIF4-dependent thermomorphogenesis in 25-day-old adult plants.

A second question was whether loss of *ELF3* function can affect thermoresponsive flowering in the Ws strain under other temperature conditions. We therefore assayed flowering in Ws and the Ws-derived null mutant *elf3-4* at 16°C and 22°C (Fig. S2B). Under these conditions, Ws robustly accelerated flowering at 22°C, whereas *elf3-4* showed no perceptible difference in flowering between the two temperatures. Thus, ELF3’s role in thermoresponsive flowering is not restricted to the Col strain or a certain temperature, but rather is necessary for whatever thermoresponsive reaction norm a strain may have for flowering.

### ELF3 and PIF4 regulate adult thermoresponsive phenotypes independently

If ELF3 and PIF4 were independent in controlling thermal responses of adult phenotypes under long days, then *elf3 pif4* double mutants would show approximately additive phenotypes. We generated *elf3 pif4* double mutants and subjected them to the same experiments as above. Our results indicated that flowering and petiole elongation constitute independent temperature responses, with ELF3 controlling the former and PIF4 controlling the latter in additive fashions (Fig. 4). That is, *elf3 pif4* double mutants showed negligible thermoresponsive flowering like *elf3*, and a negligible petiole response like *pif4*. Additionally, *elf3 pif4* flowered slightly later than *elf3* at 22°, while maintaining a negligible thermal response in flowering, indicating that *elf3* mutants are not simply restricted by a physiological limit of early flowering. The additivity of these phenotypes establishes that, under these conditions, ELF3 and PIF4 likely operate in separate thermal response pathways.

**Fig. 4.**
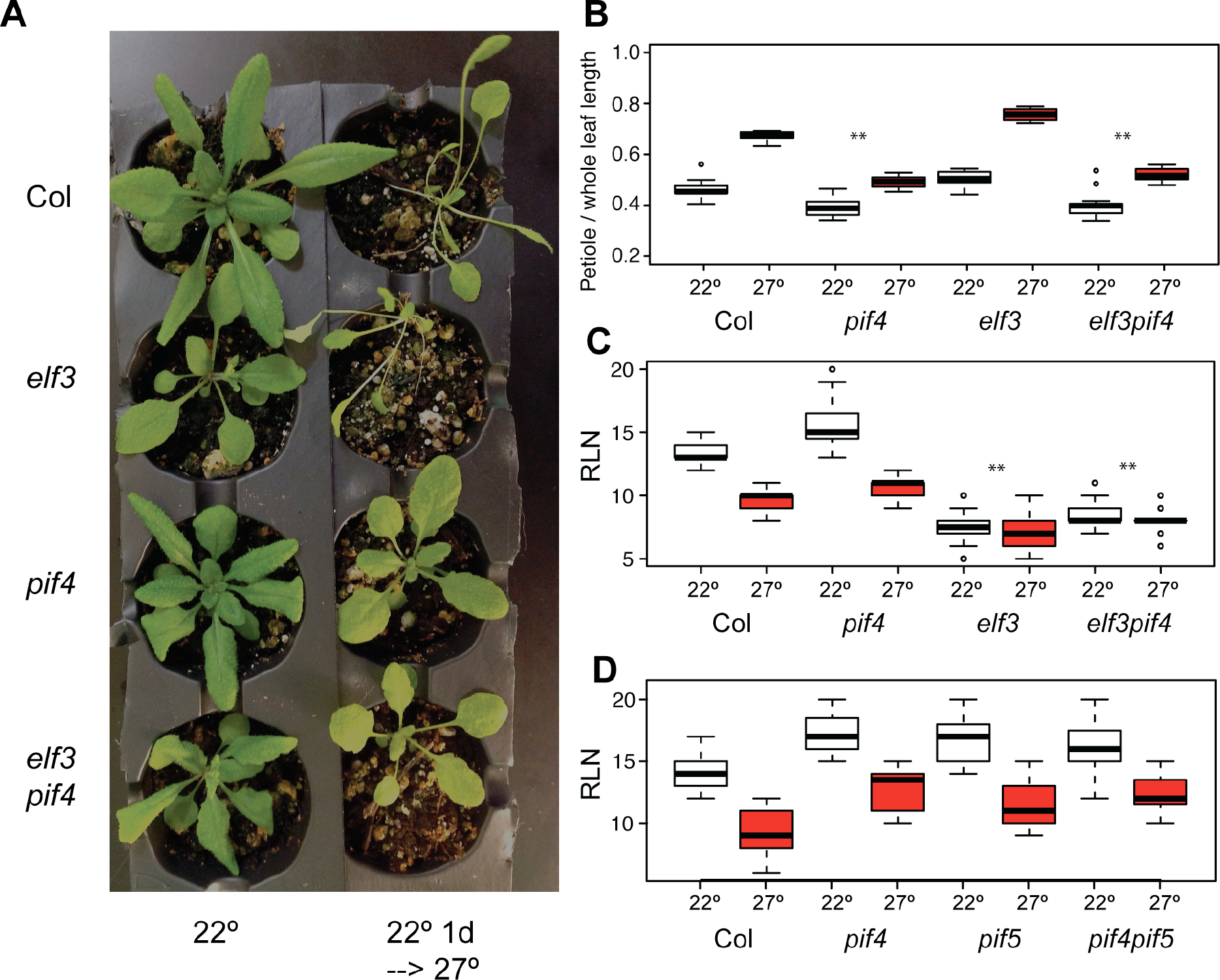
Double mutant analysis confirms PIF4 and ELF3 independence in adult temperature responses and non-redundancy of PIF4 with PIF5. (A): Col, *elf3-200*, *pif4-2*, and *elf3-200 pif4-2* plants grown under long days with two different temperature regimes were photographed at 25 days post germination. (B): Petiole elongation responses of the indicated genotypes, measured by ratio of petiole to whole leaf length at 25 days post germination. (C): Flowering temperature response of indicated genotypes, measured by rosette leaf number (RLN) at flowering. (B) and (C): n > 8 plants for each genotype in each treatment. All “27°” plants were seeded and incubated one day at 22° before transfer to 27°. Experiments were repeated with similar results. Regression analysis of data reported in Tables S6 and S7. In each case, **: Bonferroni-corrected p < 0.01, **: Bonferroni-corrected p < 0.05, .: Bonferroni-corrected p < 0.1 in testing the interaction term (different response from Col).

Previous studies have indicated that other members of the PIF family have negligible or minor (in the case of PIF5) roles in these same thermal response phenotypes [11,27,43]. For instance, under SD, *pif1 pif3 pif4 pif5* mutants behave essentially identical to *pif4 pif5* mutants in flowering response and *FT* expression, which in turn show only a very slight abrogation of these responses relative to *pif4* mutants [31]. *pif4 pif5* double mutants do show slightly abrogated thermoresponsive flowering under 12 hour light: 12 hour dark photoperiods relative to single mutants [29], similar to other thermally responsive phenotypes [11,29–31]. These previous findings, combined with the completely intact flowering response of *pif4* mutants, suggest that redundancy between PIFs plays little meaningful role in this response. However, to directly address this possibility, we evaluated thermoresponsive flowering in *pif5* and *pif4 pif5* mutants (Fig. 4D), because PIF5 is most often considered to act redundantly with PIF4 [20,29,31,44], and the only other PIF to show any small contribution to thermoresponsive flowering [29–31]. As expected, both *pif5* single mutants and *pif4 pif5* double mutants demonstrate intact thermoresponsive flowering. These observations indicate that redundancy with other PIFs is not responsible for the apparent independence of PIF4 and ELF3. Notably, petiole elongation at elevated temperatures is equally disrupted in *pif4* and *pif4 pif5* mutants, but intact in *pif5* single mutants (Figure S3), reproducing the known dependence of this trait upon PIF4 alone [11]. Consequently, our results support the previously-suggested dominance of thermomorphogenesis by PIF4 rather than other PIFs, and the irrelevance of PIF4 (and most likely other PIFs as well) to thermoresponsive flowering under LD.

Overall, the strong photoperiod-dependence of PIF4-related thermoresponsive flowering necessitates the existence of some pathway or pathways independent of PIF4 under long days, given the persistence of the phenomenon under these conditions. Based on our data, ELF3 acts in one such pathway.

### Thermoresponsive flowering under long days can operate through the photoperiodic pathway

ELF3 operates in thermoresponsive flowering at low ambient temperatures via the photoperiodic pathway, through repressing *GI* expression, after which GI in turn directly activates FT [45,46]. To evaluate whether this pathway might explain our results, we measured transcript levels of *GI* and *CO* in wild-type and *elf3* mutants under 22°C and 27°C (Fig. 5A). We found that *GI* is strongly up-regulated in *elf3* null mutants of Col and Ws backgrounds, confirming previous reports in Col [39,46]. Further, wild-type Ws showed higher basal *GI* levels compared to Col, which did not increase at higher temperatures. In contrast, Col showed very low basal *GI* levels that increased at higher temperatures to approximately the same levels as Ws. *CO* levels, however, were not substantially increased by either *elf3* mutation or increased temperature, consistent with previous reports [8,46]. Thus, robust thermoresponsive flowering was correlated with low basal levels of *GI*, and with temperature-dependent *GI* up-regulation, as observed in Col. The ELF3-dependent thermal responsiveness of *GI* expression confirms previous reports [15], though the among-strain variation in responsiveness appears to be novel and correlated specifically with flowering induction (but not hypocotyl or petiole elongation, Figures 1 and 2). High basal *GI* levels in Ws may be associated with other thermoresponsive deficiencies at high temperatures in this strain [42,47,48]. These observations support the model under which ELF3 acts in the photoperiodic pathway to engender thermoresponsive flowering, just as it does in response to lower ambient temperatures [8,46].

We attempted to measure *FT* transcript levels in these samples, expecting that they would be elevated in the early-flowering *elf3* and 27°C conditions (Figure S4). Unexpectedly, we observed that, while *FT* levels may increase slightly in the *elf3* mutants, *FT* appears dramatically down-regulated in all 27°C samples. This finding isdifficult to interpret in light of the phenotypic data, as most models of thermoresponsive flowering agree that signaling operates through *FT* [7–9,29–31], suggesting rather that these 7-day-old seedlings may be too young to measure physiologically relevant *FT* expression, at least under 27°C conditions.

**Fig. 5.**
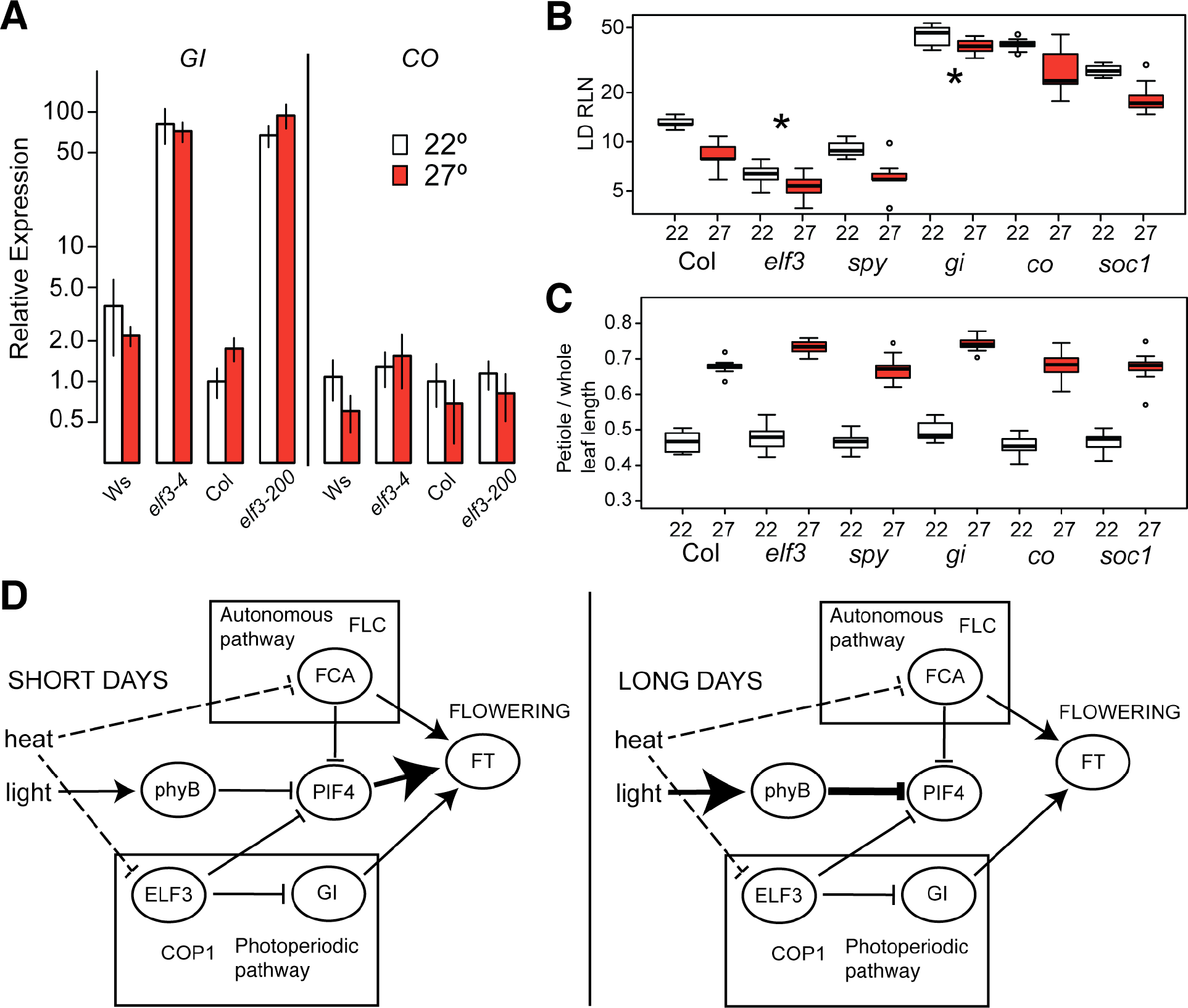
ELF3 and GI regulate thermoresponsive flowering. (A): Temperature-responsive expression of photoperiodic pathway components at ZT0. Expression of each gene is quantified relative to levels in Col at 22° (Col 22 = 1.0). Error bars represent SEM across three biological replicates. *elf3-4*: *elf3* null in Ws background; *elf3-200*: *elf3* null in Col background. (B): Thermoresponsive flowering in various flowering mutants. LD RLN = rosette leaf number at flowering under long days. *: interaction term for genotype by environment at Bonferroni-corrected p < 0.05; details of regression model in Table S9. (C) Thermoresponsive petiole elongation in various flowering mutants. For (B) and (C), n >= 8 plants of each genotype in each condition; white boxes indicate measurements at 22°, red boxes indicate measurements at 27°. *gi: gi-2, co: co-101, spy: spy-3, soc1: soc1* T-DNA insertion, *elf3: elf3-200*.This experiment was repeated with similar results. (D): Models of thermoresponsive flowering under long and short photoperiods. Dashed edges indicate speculated temperature sensing mechanisms. Edges with increased weight indicate relative increases of influence between conditions. Pathways are indicated, along with other important actors reported elsewhere.

If the photoperiodic pathway contributes to thermoresponsive flowering at elevated ambient temperatures in long days (LD), we would expect mutants in this pathway to show abrogated thermal responses, as they do under short days (SD), along with members of the autonomous pathway [7]. These two pathways also contribute independently to thermoresponsive flowering at low temperatures (16°C vs. 23°C) [6, 8]. Altogether, we would expect that a photoperiodic thermoresponsive flowering pathway would operate independently of both PIF4 and the autonomous pathways in long days. It is not clear whether the autonomous pathway would be independent of PIF4, given known interactions between FCA and PIF4 [13].

To evaluate whether these past results under other conditions also apply to long days and elevated temperatures, we measured flowering time at 22°C and 27°C in mutants in the photoperiodic pathway (*gi*, *co*, Fig. 5B). We also tested mutants of the gibberellin pathway (*spy*), and a terminal floral integrator (*soc1*), which we do not expect to be necessary for thermoresponsive flowering. We found robust thermal responses in all mutants except *elf3* and *gi*, similar to previous results under different conditions [7,8,45,46]. These results emphasize once again that differences in thermoresponsive flowering are not generalizable between photoperiods, as it has recently been shown that *co* mutants have a partial flowering acceleration defect under SD. These results implicate GI (but not CO) as an actor in thermoresponsive flowering at elevated temperatures. Collectively, these experiments suggest that the photoperiod pathway is necessary in promoting thermoresponsive flowering in long days, and expression data in this and other studies suggests that ELF3 is likely to act within this pathway.

## DISCUSSION

ELF3 and PIF4 are both crucial integrators of temperature and light signaling in controlling *A. thaliana* development. Recent literature has emphasized the centrality of PIF4-dependent thermoresponsive regulation in a variety of phenotypes, including in flowering [9,10,32]. Here, we show that PIF4 is dispensable for thermoresponsive flowering under long photoperiod conditions [11], and that ELF3 is essential for thermoresponsive flowering under these conditions. Our results integrate previous knowledge about thermoresponsive flowering, and identify at least one pathway for this response that does not involve PIF4. Moreover, we show that while polyQ variation in ELF3 affects ELF3 function, the polyQ tract is unlikely a temperature-responsive component in itself. Our results allow us to integrate the many disparate findings of current studies into classic models of thermal responses in *A. thaliana*, allowing a comprehensive view of the genetic underpinnings of this agronomically crucial plant trait.

### ELF3 polyglutamine variation appears to affect thermoresponsive traits by modulating overall ELF3 activity

In previous work, we demonstrated that polyQ variation in ELF3 is (i) common, (ii) affects many known ELF3-dependent phenotypes, and (iii) is dependent on the genetic background [35]. Following the recent discoveries that ELF3 is involved with thermal response [14–16], we confirmed that ELF3 polyQ variation also affects thermal response phenotypes in a background-dependent fashion. However, we found little support for the hypothesis that the polyQ tract has a special role in temperature sensing. Instead, as was the case for other ELF3-dependent phenotypes, ELF3 polyQ variation appeared to affect overall ELF3 functionality, with less functional ELF3 variants lacking robust temperature responses. However, a more exhaustive series of polyQ variants may be required for revealing polyQ-specific effects, in particular because the molecular mechanism(s) by which polyQ variation affects ELF3 functionality remain unknown.

### ELF3-PIF4 relationship in thermomorphogenesis

One question that remains unanswered is to what extent ELF3 participates in PIF4-dependent thermoresponsive morphologies. While our study and previous work [14,16,39] support a PIF4-ELF3 link in thermoresponsive hypocotyl elongation, this relationship disappears in the analogous case of thermoresponsive petiole elongation. These results can be explained by many hypotheses. For instance, it is possible that ELF3 regulation of PIF4 is only relevant at the early seedling stage. Another possible hypothesis is that ELF3 regulation of PIF4 in some instances is sufficient but not necessary for thermal responses. More studies are needed to understand the mechanistic details of the ELF3 and PIF4 relationship in thermomorphogenesis.

### Natural variation in temperature response

Several studies have found that different *A. thaliana* strains respond to temperature differently, either shifting or inverting the reaction norm of the phenotype in question [42,47,48]. Ws has a shifted reaction norm with respect to temperature compared to Col for photoperiod-related phenotypes, including flowering. For instance, Ws displays accelerated flowering at 23°C vs. 16°C [42], but accelerates flowering no further at 27°C. Here, we show that this acceleration requires ELF3, like the elevated temperature acceleration in Col. Another example of differential mutational effects among strains is that *gi* mutants in the *Ler* background display robust thermoresponsive flowering [6,7]. It is unclear whether this finding is due to altered wiring of pathways between these backgrounds.

### Thermoresponsive flowering requires either PIF4 or ELF3, depending on photoperiod

Under various conditions, both ELF3 and PIF4 have been found to be crucial for thermoresponsive flowering. Other members of the autonomous and the photoperiodic pathways have also been implicated in thermoresponsive flowering [6–8] (besides other pathways, [49]). Consequently, some combination of these pathways, modulated by experimental conditions, must require ELF3 and/or PIF4. We and others [11,29] have observed that PIF4 and its paralogs are not required for proper thermoresponsive flowering under longer photoperiods. Furthermore, we and others [8,46] have shown that ELF3 and the photoperiod pathway (excluding CO) are essential for proper thermoresponsive flowering under long days. It has been previously shown that PIF4 and the photoperiodic pathway contribute to thermoresponsive flowering via independent pathways [9], suggesting that under longer photoperiods PIF4 activity is inhibited, allowing other mechanisms to dominate thermoresponsive flowering.

We propose a model of thermoresponsive flowering, in which PIF4, ELF3, the photoperiodic pathway, and other pathways interact depending upon condition and genetic background (Fig. 5D). Under short days or other short photoperiods, phyB activity is down-regulated, leading to up-regulation of PIF4 [22,50–52], which at high levels occupies the promoter of the flowering integrator *FT* and induces flowering [9]. However, under longer photoperiods, phyB up-regulation leads to an attenuation of PIF4 activity, and consequently the role of PIF4 and other PIFs becomes negligible [11]. This allows canonical ambient temperature responses (such as the photoperiodic pathway, including ELF3, [8,46]) to take a dominant role in thermoresponsive flowering. Constitutive overexpression of PIF4, PIF5, and PIF3 under long day conditions induces early flowering [30], supporting the hypothesis that differences in PIF levels underlie the photoperiod-dependence of PIF4’s role. We have not formally excluded the possibility that members of the large PIF family other than PIF4 and PIF5 might contribute to the phenotype; however, there is no evidence at present to suggest that they might [11,27,30,31]. Several reports have indicated that GI and COP1, but not CO, are involved in thermoresponsive flowering [7,8,46], with GI directly binding the *FT* promoter [46]. Under each of these conditions, FT-induced flowering is activated by a different signaling cascade. This interpretation leads to a coherent view of how light and temperature responses are integrated in this important plant trait.

To summarize, at least three independent mechanisms have been described that promote thermoresponsive flowering in any context. These include the photoperiodic pathway (PHYB/ELF3/GI/COP1), the autonomous pathway (PHYA/FCA/FVE/TFL1/FLC), and the PIF4-dependent pathway (PIF4/H2A.Z/gibberellin), all of which converge by regulating *FT* (although the last pathway may also act through other integrators [29,30]). Our results showing a down-regulation of *FT* at elevated temperatures in young seedlings may also hold clues regarding the interaction of these pathways, but are difficult to interpret in the context of the other evidence. The collective results of our experiments and previous work suggest that the first two pathways are necessary but not sufficient for thermoresponsive flowering, and that the third (PIF4) is sufficient but not necessary for thermoresponsive flowering. Further study will be necessary in understanding the interdependencies of the three pathways. For instance, it has been suggested that PIF4 binding to the *FT* promoter is dependent on cooperativity with a second photoperiod-controlled actor [34].

In conclusion, we observe that ELF3 is involved in the hypocotyl response to elevated temperature as reported previously, and that this response can be abrogated by poorly-functioning ELF3 polyQ variants. We further demonstrate that ELF3 has little effect on the petiole temperature response, and is necessary for the flowering temperature response, suggesting that it functions independently of PIF4, potentially in the photoperiodic pathway. These results reiterate the complexity of these crucial environmental responses in plants, and can serve as a basis for further development of our understanding of how plants respond to elevated temperatures. In the context of climatic changes, this understanding will serve those attempting to secure the global food supply.

## MATERIALS AND METHODS

*Plant materials and growth conditions*. All mutant lines (except *pif4-2 elf3-200*) were either described previously or obtained as T-DNA insertions from the Arabidopsis Biological Resources Center at Ohio State University [53,54],and are described in Table S11. *pif4-2 elf3-200* was obtained via crossing and genotyping. T-DNA insertions were confirmed with primers described in Table S10. For hypocotyl assays, seedlings were grown for 15d in incubators set to SD (8h light: 16h dark days, with light supplied at 100 μmol. m^−2^s^−1^ by cool white fluorescent bulbs) on vertical plates as described previously [35]. All plates were incubated at 22° for one day, after which one replicate arm was transferred to an incubator set to 27°, with another replicate arm maintained at 22°. For flowering time assays, plants were stratified 3-5d at 4° in 0.1% agarose and seeded into Sunshine #4 soil in 36-pot or 72-pot flats to germinate at 22° under LD (16h light: 8 hr dark days, with light supplied at 100 μmol.m^−2^s^−1^ by cool white fluorescent bulbs). Replicate arms were subsequently transferred to 27° LD conditions as indicated, with others remaining at 22°. Different temperature treatments of the same experiment were identical with respect to randomization, setup, and format. At 25d, petiole length and whole leaf length (including petiole) of the third leaf were measured, and the ratio of these values was further analyzed. Flowering was defined as an inflorescence ≥1cm tall; at this point, date and rosette leaf number were recorded.

*Trait data analysis*. All data analysis was performed using R v3.2.1 [55]. Where indicated, temperature responses were modeled using multiple regression in the form *Phenotype* **~ μ +β_G_***Genotype* **+ β_T_***Temperature* **+ β_G×T_**(*Genotype* × *Temperature*) **+β_E_***Experiment* + *Error*. All experiments were included in models for transgenic experiments, and thus the **β_E_** term describes systematic variation between experiments, whereas line-specific effects among transgenics should be modeled in the error term. Where temperature responses are reported, they consist of the **βT + βG×T** terms and associated errors 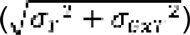 where **σ_T_** is the standard error for **β_T_** and **σ_G×T_** is the standard error for **β_G×T_**), and thus are corrected for systematic experimental variation and temperature-independent genotype effects. Where p-values are reported for G×T interaction effects, they have been subjected to a Bonferroni correction to adjust for multiple comparisons. Analysis scripts and data are provided at https://figshare.com/articles/elf3pif4datacodev2/3398353.

*Gene expression analyses*. Seedlings were grown for 1d under LD at 22°, after which one replicate arm was transferred to LD at 27°, with another replicate arm remaining at 22°, and all seedlings were harvested 6d later at indicated times. At harvest, ~30mg aerial tissue of pooled seedlings was flash-frozen immediately in liquid nitrogen and stored at −80°. RNA extraction, cDNA synthesis, and real-time quantitative PCR were performed as described previously [35], using primers in Table S10. Transcript levels were quantified from technical triplicate across at least two biological replicates using the **ΔΔ**C_t_ method, assuming 100% primer efficiency [56].

## ACKNOWLEDGMENTS

We thank Philip Wigge and Jaehoon Jung for ideas, helpful conversations, sharing unpublished data, and comments on this manuscript. We thank Evan Eichler for use of the LightCycler instrument. We thank members of the Queitsch lab for helpful discussions. This work was supported by National Institutes of Health New Innovator Award DP2OD008371 to CQ.

## Supporting Information Captions

**Fig. S1. Expression analysis of *PIF4* and *AtHB2* depends on temperature, genetic background, and ELF3 functionality.** Error bars represent the standard deviation across two biological replicates. White bars represent 22° expression, red bars 27° expression for each line. Tissue was collected from 7d seedlings at ZT0.

**Fig. S2. Regulation of adult thermoresponsive traits by ELF3 and FCA is independent of PIF4 and modulated by genetic background.** Flowering temperature response of indicated genotypes under indicated conditions, measured by petiole length to leaf length ratio at 25 days or rosette leaf number (RLN) at flowering. For each experiment, n > 10 plants for each genotype in each treatment. Regression analysis of data in Tables S4 and S5.

**Fig. S3. Regulation of adult thermoresponsive petiole elongation traits occurs principally through PIF4.** Petiole elongation temperature response of indicated genotypes under indicated conditions, measured by ratio of petiole length to leaf length at 25d. For each experiment, n > 10 plants for each genotype in each treatment. This experiment was repeated with similar results. Regression analysis of data in Table S8.

**Fig. S4. Expression of *FT* in 7d seedlings responds to temperature and *elf3* status.** White bars represent 22° expression, red bars 27° expression for each line. Tissue was collected from 7d seedlings at ZT0. Error bars indicate SEM across three biological replicates.

**Table S1. Regression analysis of hypocotyl elongation temperature response among Col and Ws transgenic lines.**

**Table S2. Regression analysis of petiole: leaf length ratio and rosette leaf number at flowering temperature response among Col and Ws transgenic lines.**

**Table S3. Regression analysis of rosette leaf number at flowering and petiole: leaf length ratio temperature responses in *elf3* and *pif4*.**

**Table S4. Regression analysis of rosette leaf number at flowering temperature response in Ws and *elf3-4*.**

**Table S5. Regression analysis of petiole: leaf length ratio temperature response in Col and *fca* mutants.**

**Table S6. Regression analysis of rosette leaf number at flowering temperature response in *elf3 pif4* double mutants.**

**Table S7. Regression analysis of rosette leaf number at flowering temperature response in *pif4 pif5* double mutants.**

**Table S8. Regression analysis of the petiole elongation temperature response in *pif4 pif5* double mutants.**

**Table S9. Regression analysis of rosette leaf number at flowering and petiole: leaf length ratio temperature responses in flowering pathway mutants.**

**Table S10. Primers used in this study.**

**Table S11. Mutant lines used in this study.**

